# Strain- or Stress-sensing in mechanochemical patterning by the phytohormone auxin

**DOI:** 10.1101/582551

**Authors:** Jean-Daniel Julien, Alain Pumir, Arezki Boudaoud

**Affiliations:** Reproduction et Développement des Plantes, Université de Lyon, ENS de Lyon, UCB Lyon 1, CNRS, INRA, Lyon Cedex 07, France; Laboratoire de Physique, Université de Lyon, ENS de Lyon, UCB Lyon 1, CNRS, Lyon Cedex 07, France; Max-Planck Institute for Dynamics and Self-Organisation, Göttingen, D-37077, Germany

**Keywords:** patterning, chemomechanical model, auxin transport, shoot apical meristem

## Abstract

Both chemical and mechanical fields are known to play a major role in morphogenesis. In plants, the phytohormone auxin and its directional transport are essential for the formation of robust patterns of organs, such as flowers or leaves, known as phyllotactic patterns. The transport of auxin was recently shown to be affected by mechanical signals, and conversely, auxin accumulation in incipient organs affects the mechanical properties of the cells. The precise interaction between mechanical fields and auxin transport, however, is poorly understood. In particular, it is unknown whether transport is sensitive to the strain or to the stress exerted on a given cell. Here, we investigate the nature of this coupling with the help of theoretical models. Namely, we introduce the effects of either mechanical stress or mechanical strain in a model of auxin transport, and compare the patterns predicted with available experimental results, in which the tissue is perturbed by ablations, chemical treatments, or genetic manipulations. We also study the robustness of the patterning mechanism to noise and investigate the effect of a shock that changes abruptly its parameters. Although the model predictions with the two different feedbacks are often indistinguishable, the strain-feedback seems to better agree with some of the experiments. The computational modeling approach used here, which enables us to distinguish between several possible mechanical feedbacks, offers promising perspectives to elucidate the role of mechanics in tissue development, and may help providing insight into the underlying molecular mechanisms.

## 1 Introduction

Understanding the formation of patterns in living organisms has long intrigued scientists [1]. This phenomenon has been investigated in a wide variety of species, such as zebrafish [23] or hydra [4]. We focus here on patterning in plants. The highly regular positioning of the organs in plant shoots, called phyllotaxis, relies on the patterns of the phytohormone auxin, whose local accumulation is necessary for the emergence of new organs [24, 18]. Auxin efflux is facilitated by the membrane-localized PIN FORMED1 (PIN1) protein [35, 51], which can be polarly (asymmetrically) distributed, inducing a directional transport of auxin, and resulting in well-defined auxin patterns. This transport is essential for the development of the plant. Indeed, knocking down the transporter activity, either genetically or chemically, leads to the absence of organ primordia at the shoot tip. Organ development can be then rescued by local application of exogenous auxin [34, 40, 41].

Several models have been proposed to describe the interaction between the hormone and its carrier in the context of the shoot apex, with the major assumption that the transporters allocation between the different sides of a cell is driven by auxin fluxes [49] or by auxin concentrations [48, 22, 44, 47]. Here we focus on the second assumption, which has received more attention, because how the auxin concentration regulates the transporters remains unclear.

Since morphogenesis relies on changes in the structural elements of the organism, biochemical patterns must also influence the mechanics of these structural elements, in plants [16] and in animals [19, 9, 39]. Conversely, mechanics can feedback on biochemical processes [16, 19, 20, 45], including gene expression [11] and cell fate [3]. Plants are well suited to study the coupling between biochemical and biophysical processes as hydrodynamic cell pressure generates tremendous forces and results in a high tension in the polysaccharide-based walls surrounding cells [13, 16]. Recent progress in computational approaches [8] and in the measurements of cell mechanics [28] has fostered a renewed interest in the mechanics of plant morphogenesis.

Notably, at the shoot apex, organogenesis is associated with a decrease in the stiffness of the cell wall [36] and likely with a reduced mechanical anisotropy [46]. In the context of phyllotaxis, the coupling between mechanics and chemistry was explored theoretically [33] and mechanics has been proposed to regulate the transport of auxin in the shoot apical meristem [17]. A mechanical feedback was postulated, that was capable of generating a local accumulation of auxin [17]. Based on preferential localization of PIN1 in the regions of the plasma membrane in contact with the wall with highest mechanical stress, this mechanical feedback is supported by several experiments [17, 32, 7]. However, whether PIN1 polarity is driven by the stress or by the strain of the cell wall remains unclear. More generally, the issue of stress- or strain-sensing calls for further investigation, especially in the context of plant development [31].

Here we address this issue by investigating a model of mechanical feedback on auxin transport, whose predictions can be compared with experimental results. We also study the robustness of the pattern formation mechanism with respect to noise and abrupt changes in some of the biochemical parameters. The predictions of the two mechanisms do not differ much. A feedback based on strain nonetheless leads to more realistic predictions, and to a patterning mechanism which is less sensitive to noisy inputs or sharp perturbations.

## 2 Results

### 2.1 A mechanochemical model of auxin patterning

We built a mechanochemical model of auxin transport and tissue mechanics that incorporates the influence of auxin on the stiffness of the cell wall and the mechanical feedback from the cell wall on auxin transport, as illustrated in Fig. 1A. We briefly describe its main ingredients here, deferring further details to the section ?Model formulation? below. We model a single cell layer, accounting for the epidermis of the shoot apex. We simplify tissue topology and assume the cells to form a regular hexagonal lattice, following [17], because we aim at stereotypic simulation results that would ease the comparison between strain- and stress-sensing mechanisms. The tissue is under global isotropic tension from turgor pressure, the cellular inner pressure that is assumed to be uniform. Each edge of the lattice is made of two contiguous cell walls associated with either of the two neighboring cells. Each cell wall is modeled by a linear spring, whose stiffness decreases with auxin concentration in the corresponding cell. This accounts for the positive effect of auxin on growth in the shoot apex. Changes in auxin concentration are due to production, degradation, diffusion and transport. We considered two hypotheses for the mechanical feedback: PIN1 transporters are inserted preferentially facing the cell walls that (i) undergo the highest elastic strain (elastic deformation) or (ii) the highest stress (force).

**Fig. 1.**
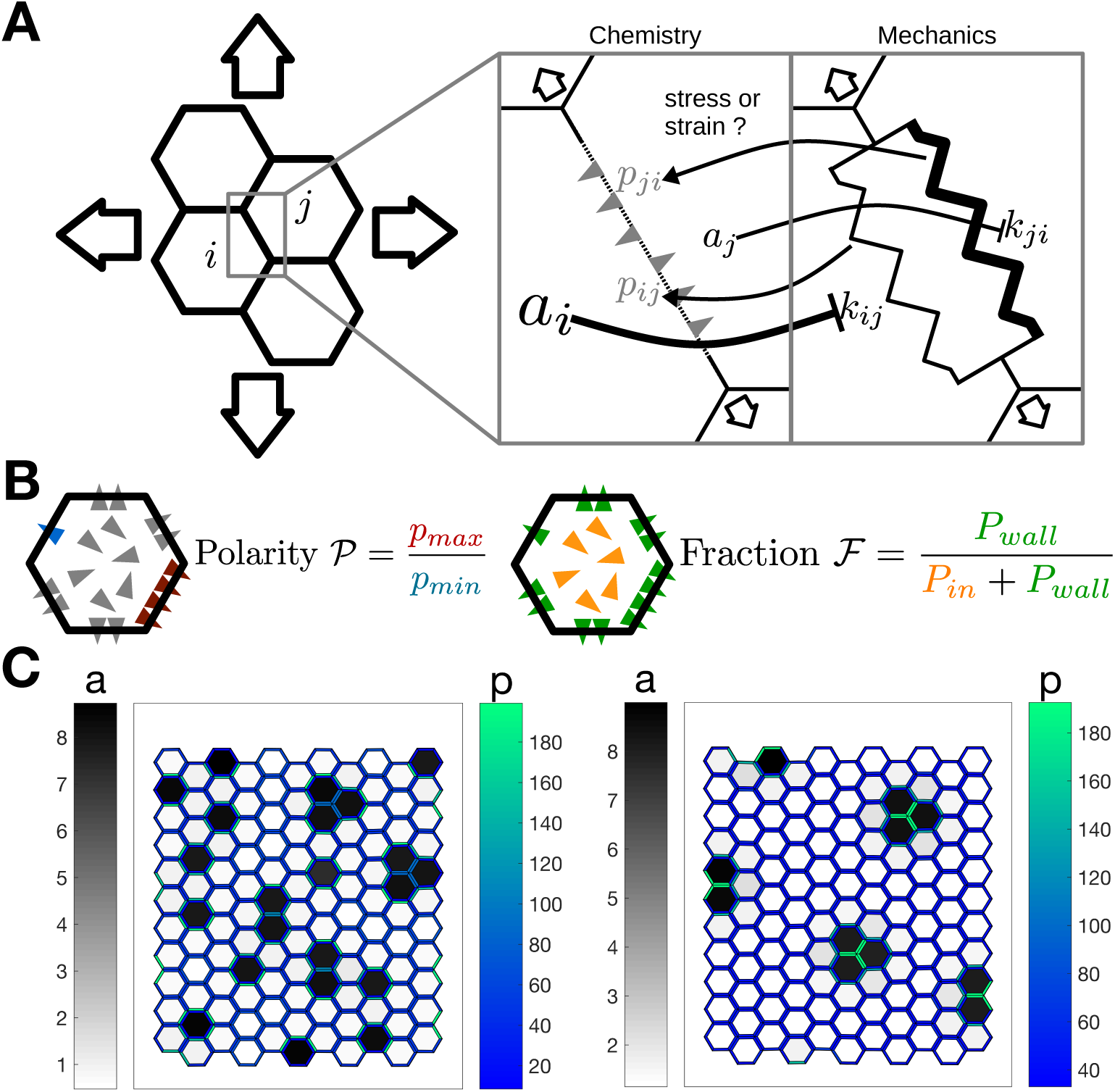
Modeling assumptions and observables: (A) Schematic of the model: the tissue consists of hexagonal cells. The auxin transporters are localized at the plasma membranes and are influenced by the mechanical status of the corresponding cells walls. The biochemical and mechanical effects are presented in two different boxes, and their interactions are represented by arrows. The tissue is stretched by the turgor pressure. PIN1 proteins facilitate auxin movement out of cells. The stiffness of each cell wall is a decreasing function of the auxin concentration in the cell. The amount of effective transporters increases with the stress or strain at the cell wall. (B) Schematic definition of the two observables: the triangles represents the PIN1 transporters, colored according to the definition of the observables. The polarity 𝒫 is the ratio between the two extremal concentrations in the cell walls of a cell. The fraction ℱ is the ratio between the number of effective transporters and the total number of transporters. (C) An example of pattern predicted by the model, with each feedback (left: stress; right:strain). The cells are colored according to their auxin concentration (gray levels), the cell walls according to the density of PIN1 transporters (blue-green levels).

We used two main observables: the polarity 𝒫, defined as the ratio between the highest and the lowest transporter concentrations along the cell edges, and the membrane fraction ℱ, defined as the ratio between the number of transporters localized at the plasma membrane and the total amount of transporters in a cell (Fig. 1B). Typical patterns obtained by solving the models numerically are shown in Fig. 1C. In order to verify our simulations, we performed an analytical linear stability analysis of the homogeneous state (supplementary text and Fig. S1) and predicted wavelengths in agreement with numerical results for a broad range of parameter values.

For a given set of parameters, wavelengths are larger for strain- than for stress-sensing (in agreement with linear stability analysis, see the supplementary note and Fig. S1). Because of this discrepancy in the wavelength, all analyses performed hereafter were also carried out on 4 other sets of parameters for the stress feedback, presented in the supplementary material (Fig. S2-6). These additional simulations largely support the conclusions presented here.

### 2.2 Cell ablation induces radial polarity

When a single epidermal cell is laser-ablated at the shoot apex, the hypothesis that the epidermis is in tension implies that the maximal stress orientation is circumferential around the ablation [15]. The preferential localization of PIN1 transporters in the stretched walls would then makes the transport radial. A model similar to ours was able to reproduce this observation with a stress-driven feedback [17]. Here, we specifically asked whether our model can, with a strain-driven feedback mechanism, reproduce the experimental results. As in [17], we modeled the ablated cell by removing auxin, its transporters, and cell walls, and by preventing the insertion of transporters in the membranes adjacent to ablated cells. Our results, see Figure 2, show a radial reorientation of the transporters in neighboring cells, as observed in experiments [17]. It thus appears that ablation experiments do not allow us to distinguish between strain- and stress-feedback mechanisms.

**Fig. 2.**
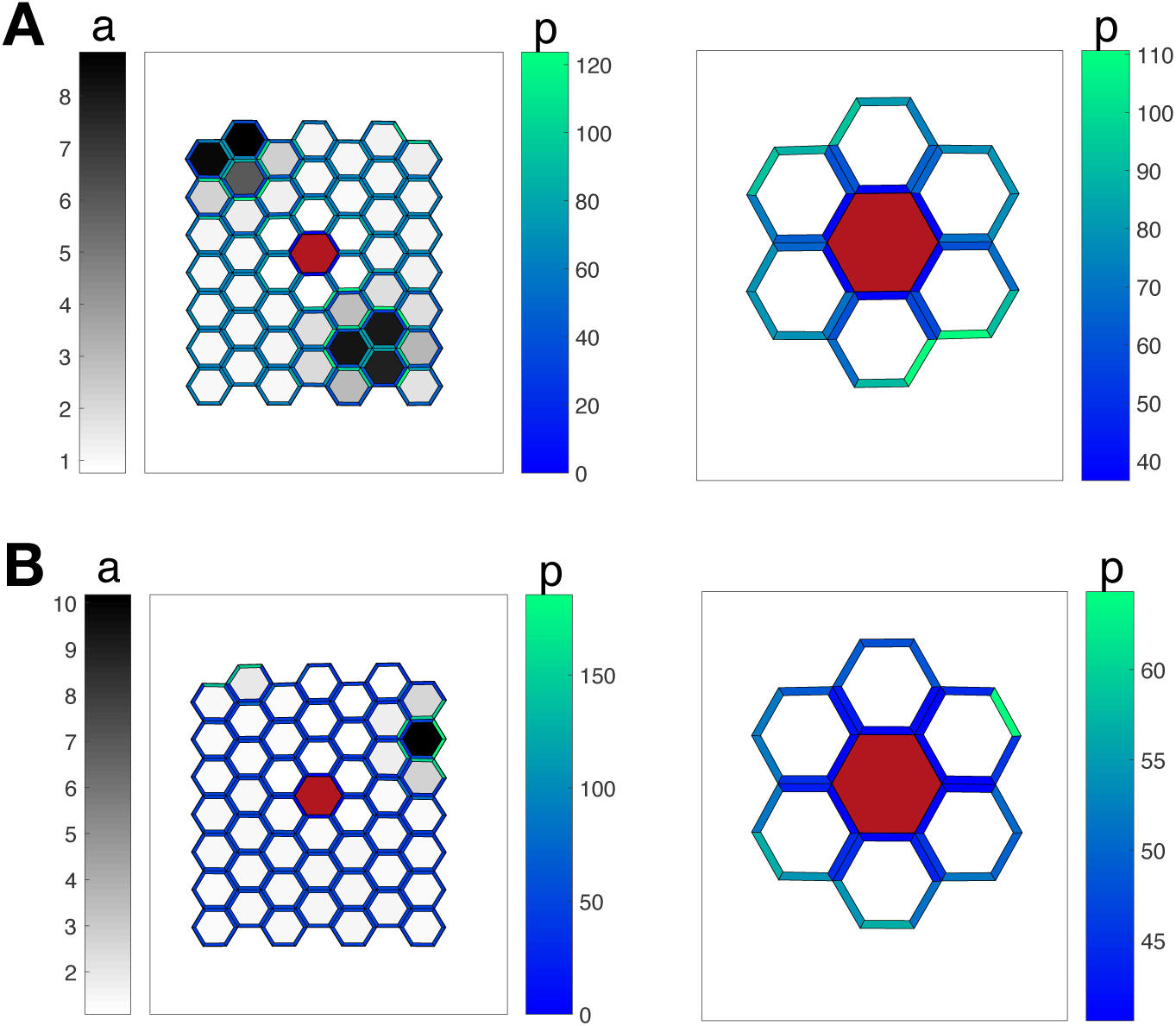
Radial polarity around a cell ablation: The red hexagon corresponds to the ablated cell, whose walls, auxin and transporters have been removed from the simulation. The transporters pointing toward this cell are also removed. The cells are colored according to their auxin concentration, the cell walls according to the concentration of transporters. The insertion of transporters is driven either by stress (A) or by strain (B). Auxin is not represented on the close-ups. Parameters for the strain feedback are indicated in Table 1. Parameters for the stress feedback are identical, except for *H*_*k*_ = 1.06, which was chosen to increase wavelength and better visualize the effect of an ablation on polarity (results are independent of the value of *H*_*k*_).

### 2.3 Variations of polarity with turgor pressure

Experiments with osmotic treatments show that PIN1 polarity 𝒫 is affected by changes in turgor pressure [32]. By performing osmotic treatments, turgor pressure was varied by approximately a factor 2 in these experiments. Polarity was significantly smaller (−15%) with reduced turgor and slightly smaller (−5%) with enhanced turgor.

**Table 1.** Values of the parameters used in the simulation: Parameters were estimated from previous models [22, 44, 17]. Stiffness dependence on auxin concentration was chosen to display 5-fold variations, in the range of realistic values [29]. The tension resulting from hydrostatic pressure was then chosen to yield a typical deformation of 4%, in the range estimated from osmotic treatments [32]. Finally, PIN1 insertion was tuned so that the fraction of transporters inserted in the membrane is around 0.7, similar to experimental estimates [32].

| $l^{(0)}$ | $\sigma$ | $s_a$ | $d_a$ | $D$ | $P$ | $K_t$ | $H_t$ | $K_f$ | $H_f$ | $k_0$ | $k_{min}$ | $\delta k$ | $K_k$ | $H_k$ |
| --- | --- | --- | --- | --- | --- | --- | --- | --- | --- | --- | --- | --- | --- | --- |
| 1 | 1.4 | 1 | 0.5 | 5 | 100 | 2 | 1 | 30 | 3 | 30 | 10 | 40 | 2 | 2 |

We simulated the effect of a gradual increase or decrease of turgor pressure by modulating tissue tension *σ* (see Eq.(1)) with respect to its reference value, *σ*_0_, and monitored the change in the polarity of the transporters with the two mechanical feedbacks. Our results, see Fig. 3, show that, irrespective of the feedback mechanism, polarity increases over a range 0.5*sigma*_0_ ≲ *σ* ≲2*σ*_0_, as observed experimentally.

**Fig. 3.**
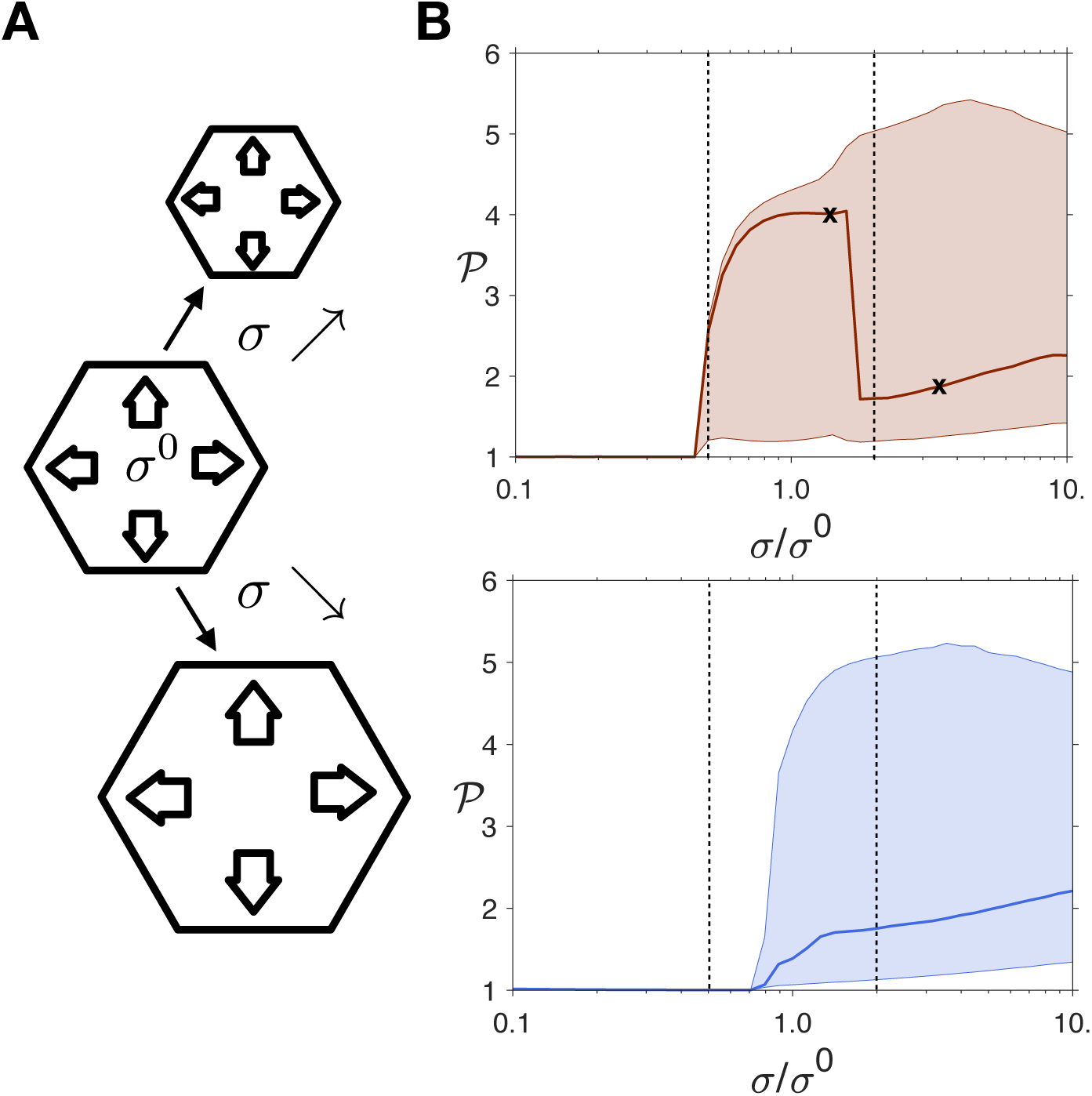
Dependence of polarity on turgor pressure: (A) Schematic of the simulations: starting from σ^(0)^, the pressure is gradually increased or decreased, for pressures ranging from 0.1 × σ^(0)^ and 10 × σ^(0)^. (B) For each cell the polarity 𝒫 is measured. The curves represent the median value of 𝒫 and the shaded areas the interval between the 15^th^ and 85^th^ percentiles in a tissue of 3600 cells. The results are shown in red (left) for the stress-based feedback, in blue (right) for the strain-based feedback. The dashed lines show the range of pressure investigated experimentally [32] and the black crosses indicate the parameters for which tissues are shown in Fig. S8.

The increase is sharper at small turgor pressure, consistent with experiments. At higher values of *σ* (*σ* ≈ 2*σ*_0_), the two models slightly disagree with experiments as they predict a slight increase in polarity while a small decrease is observed. It is unclear whether this is due to a shortcoming of the model or to a bias in experimental quantifications.

With the stress feedback, the sharp drop seen in Fig. 3 is due to a change in the overall generated pattern of auxin peaks, which is not relevant in the present context, and is followed by an increase in polarity; examples of tissues and PIN1 distributions before and after this transition are shown in Figure S2.

### 2.4 Global softening of the tissue increases polarity

Cellulose is the stiffest polysaccharide in plant cell walls. Accordingly, impairing cellulose synthesis by chemical treatments leads to the softening of plant tissues [42]. The consequence of such treatments on PIN1 localization was also investigated in [17]. First, PIN1 polarity, 𝒫, is amplified - the membrane concentration of PIN1 increases where it is high before treatment and decreases elsewhere. Second, the amount of PIN1 decreases in the cytosol, or equivalently the membrane fraction ℱ increases.

We simulated such treatments by decreasing the minimal stiffness, *k*_*min*_, of the cell walls (Fig. 4). The increase in polarity 𝒫 and in the membrane fraction ℱ is reproduced with both feedbacks (Fig. 4). Therefore, these experiments do not allow us to discriminate between the two models of feedback.

**Fig. 4.**
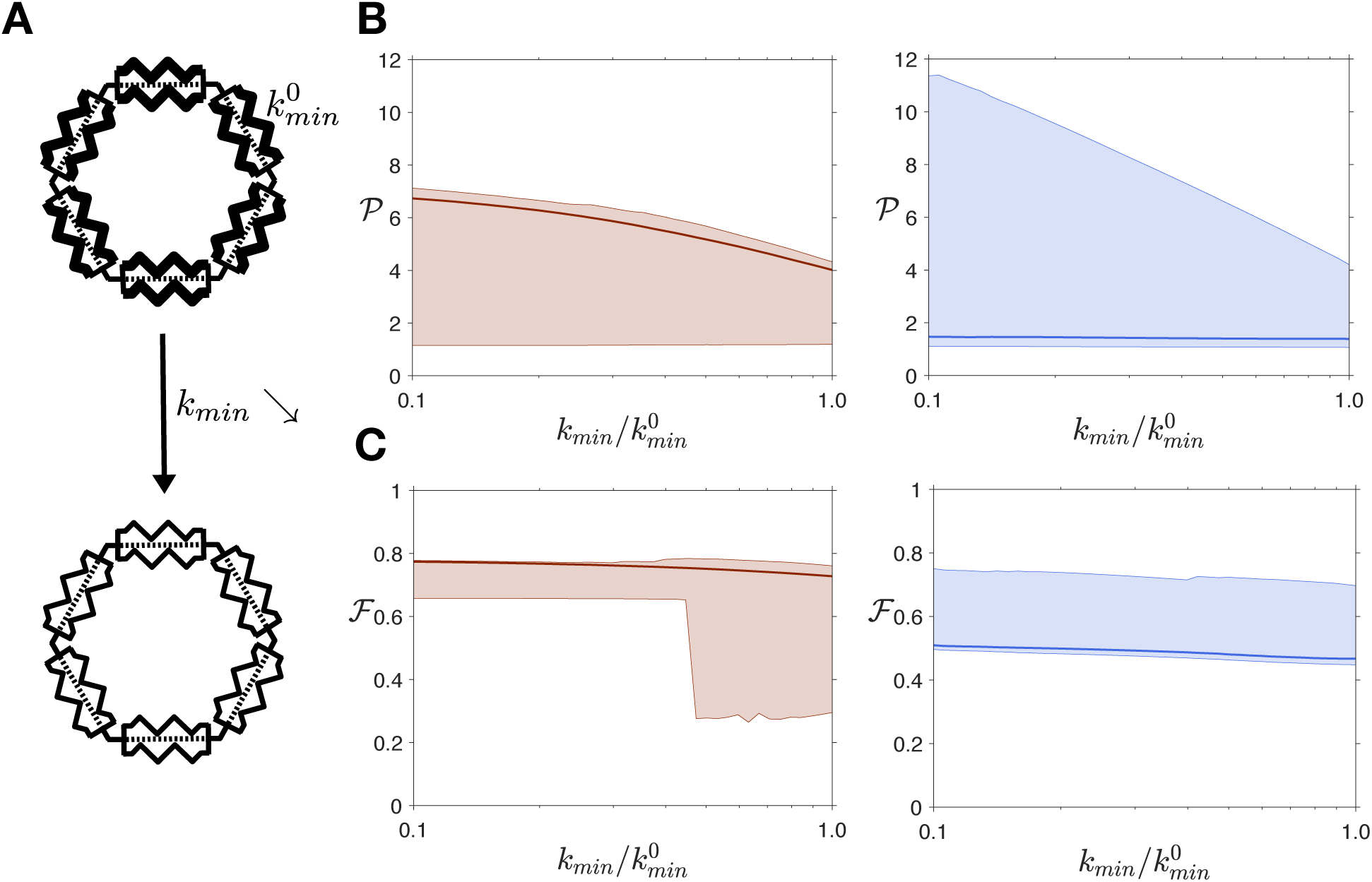
Global softening of the tissue increases polarity: (A) Schematic of the simulations: starting from 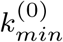, the minimal stiffness of the tissue is gradually decreased, ranging from 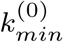 to 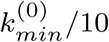. This corresponds to the stiffness at high levels of auxin. We quantified (B) the polarity 𝒫 and (C) the membrane fraction ℱ of transporters. The curves represent the median and the shaded areas the interval between the 15^th^ and 85^th^ percentiles in a tissue of 3600 cells. The results are shown in red (left) for the stress-based feedback, in blue (right) for the strain-based feedback.

### 2.5 Reducing auxin-driven softening disrupts polarity

It has been shown that de-methyl-esterification of the pectin homogalacturonan by pectin methylesterases (PMEs) is necessary for auxin to soften the cells [37]. Indeed, in plants overexpressing an inhibitor of PME activity, PME INHIBITOR3, patterns of auxin accumulation are absent and organ primordia do no form. In addition, the application of auxin does not trigger any of auxin response (no expression of the response reporter DR5), tissue softening, or organ formation [7]. In these plant lines, polarity and membrane fraction are both reduced.

To simulate such overexpression, we decreased the value of the parameter, *δk* (see Eq.(3) in the method section), that controls the sensitivity of cell wall stiffness to auxin. A vanishing *δk* means that auxin has no effect on wall stiffness, and high values of *δk* imply that a small change in auxin level induces a large change in stiffness. In parallel, we kept *k*_*min*_ + *δk* constant so that the reference stiffness in the absence of auxin remains unchanged. We measured the polarity 𝒫 and the fraction ℱ of transporters in the cell walls with the two types of feedbacks.

Fig. 5 shows that when *δk* decreases, the polarity converges towards 1, corresponding to a homogeneous state with no patterns. Thus the conclusion that the polarity is lost as a result of tissue stiffening is independent of the nature of the mechanical feedback. However, the variations of the membrane fraction ℱ depend on the precise feedback mechanism. When stiffening the tissue, ℱ decreases with the strain feedback, but remains approximately constant with the stress-feedback, as shown in Fig. 5.

**Fig. 5.**
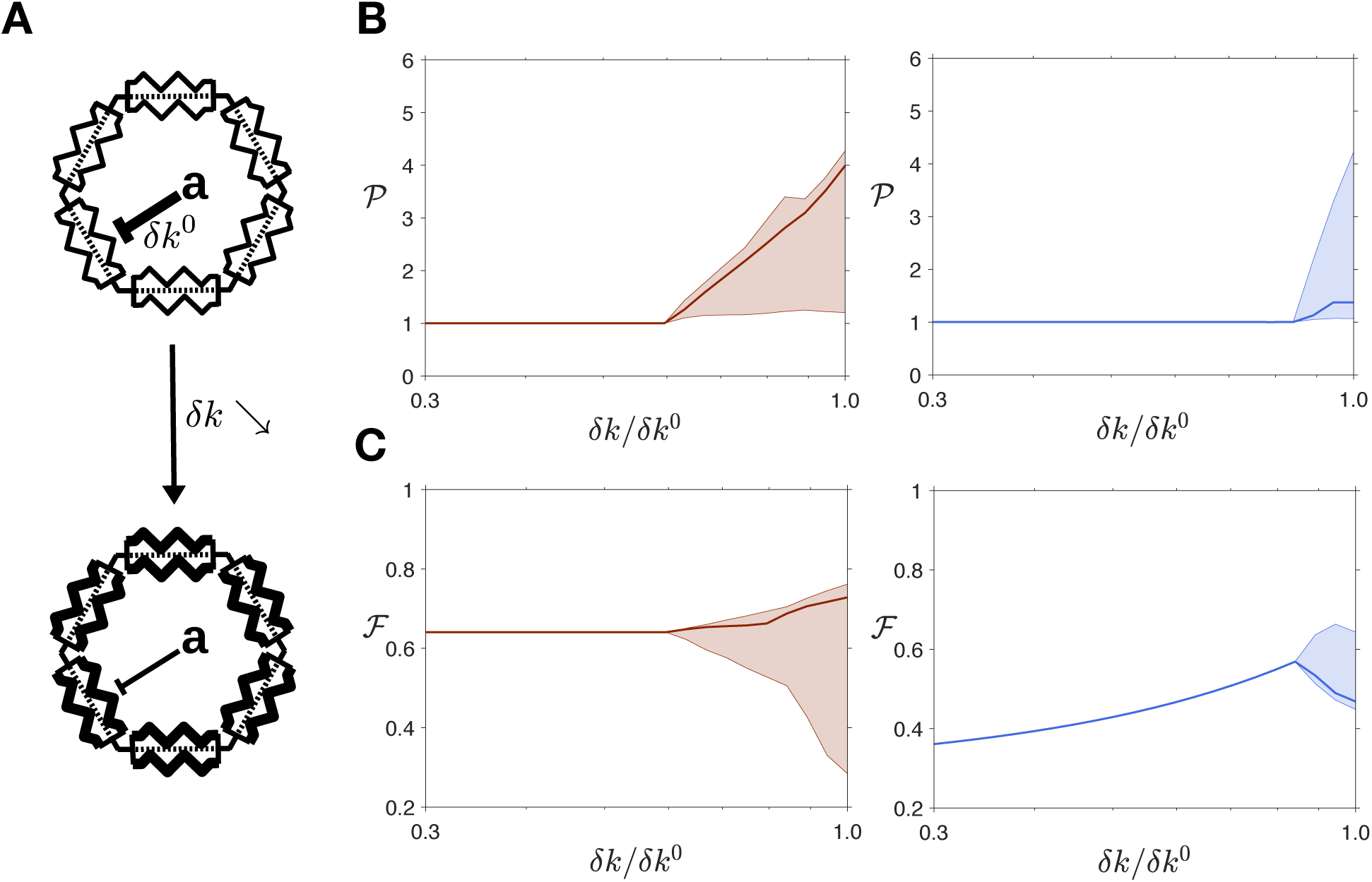
Reducing auxin effects on the cell wall disrupts polarity: (A) Schematic of the simulations: starting from *δk*^(0)^, the amplitude of stiffness variations is gradually decreased, from 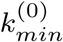 to 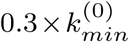. We quantified (B) the polarity 𝒫 and (C) the membrane fraction ℱ of transporters. The curves represent the median and the shaded areas the interval between the 15^th^ and 85^th^ percentiles in a tissue of 3600 cells. The results are shown in red (left) for the stress-based feedback, in blue (right) for the strain-based feedback.

The behavior of the model can be understood analytically. In the homogeneous state, that is in the absence of auxin patterns as observed in the simulations when *δk* is decreased, the membrane fraction can be computed exactly, as shown in the supplementary material. The calculations predict that ℱ^*stress*^ remains constant, whereas ℱ^*strain*^ decreases when *δk* decreases. Note that it is relevant to assume a homogeneous state of the system because naked meristems do not show auxin accumulation patterns. In view of these results, we conclude that a strain feedback is more realistic to describe experiments where PME activity is reduced.

### 2.6 Correlation between auxin level and polarity

Experiments showed that, at the initiation of a primordia, the concentration of PIN1 transporters in the plasma membrane increases with the concentration of auxin [18], likely independently of the increase in PIN1 transcription in response to auxin. Comparison of auxin and PIN1 concentrations in our simulations show that with the strain-sensing mechanism, the model qualitatively reproduces the experimental observations, whereas a stress-sensing mechanism leads to the opposite behavior, see Fig. 6. Again, analytical calculations in the limit of small auxin fluctuations around the homogeneous state confirm this result, regardless of the choice of parameters (see supplementary material). Accordingly, these results support a strain-sensing mechanism.

**Fig. 6.**
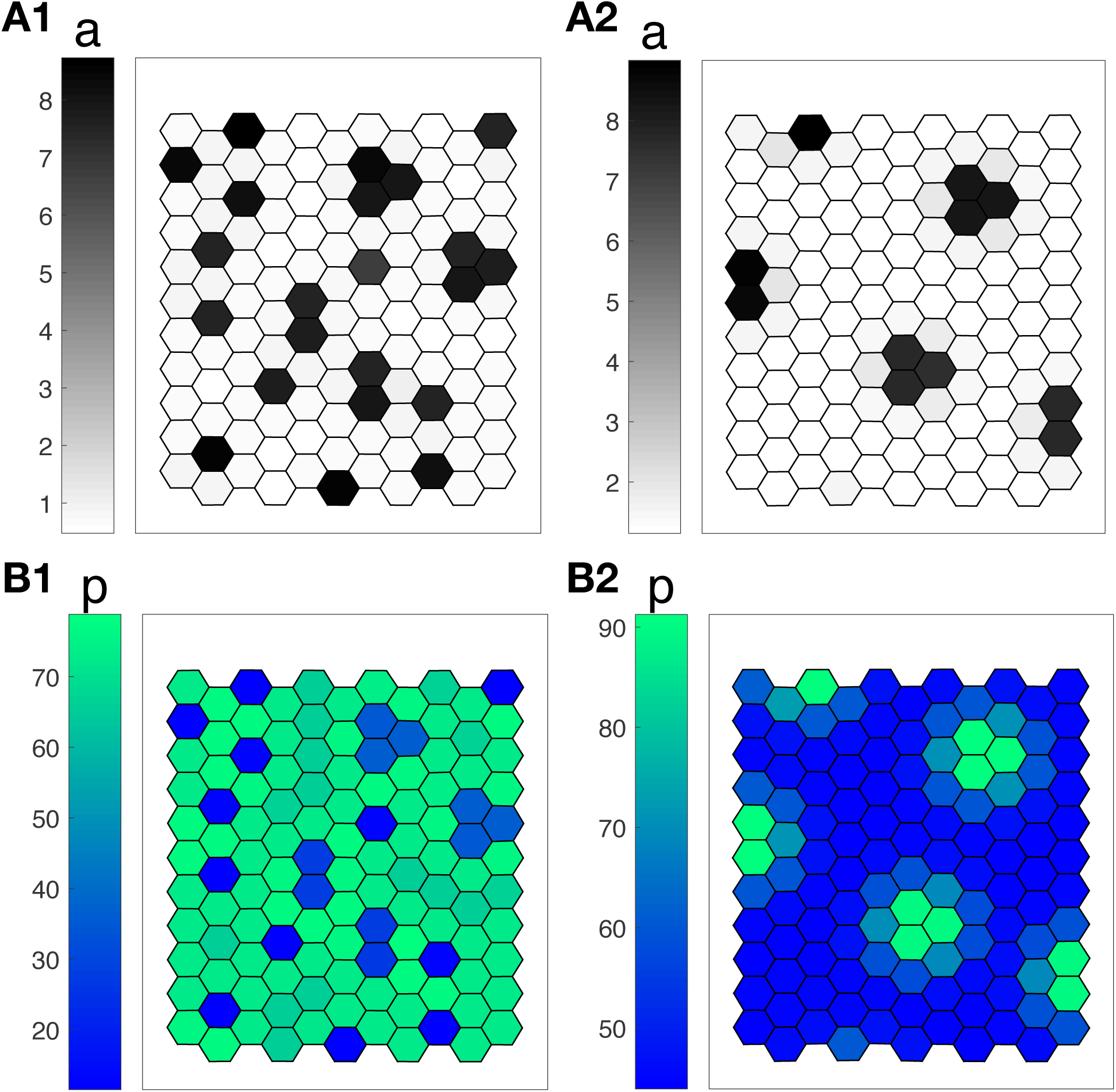
Correlation between auxin and PIN1 concentrations: Examples of pattern predicted by the model, with each feedback (A/B1: stress; A/B2: strain). The cells are colored according to their auxin concentration (gray levels, A1/2), or according to the density of PIN1 transporters, averaged over their walls (blue-green levels, B1/2).

### 2.7 Robustness to noise

We now investigate the sensitivity of the pattern to noise. To this end, a random term, temporally and spatially uncorrelated, is added to the production rate of auxin or to the concentration of transporters, respectively *s*_*a*_ or *P* in Eq. (2). We measured the wavelength of the pattern and its dependence on the noise amplitude.

As shown by panel A in Fig. 7, the pattern is almost insensitive to the effect of noise on the production of auxin. In contrast, see Fig. 7B, noise in the concentration of transporters *P* alters significantly the pattern. With the stress feedback, the wavelength is observed to increase by up to 200%, whereas the increase in the wavelength of the pattern is only 20% with the strain feedback. This difference in the sensitivity of the patterns to noise in the transporters, PIN1, also discriminates between the two models. Although it is difficult to assess the level of noise in plants, the striking robustness of phyllotaxis to fluctuations in auxin levels [50] suggests that the feedback based on strain is more plausible.

**Fig. 7.**
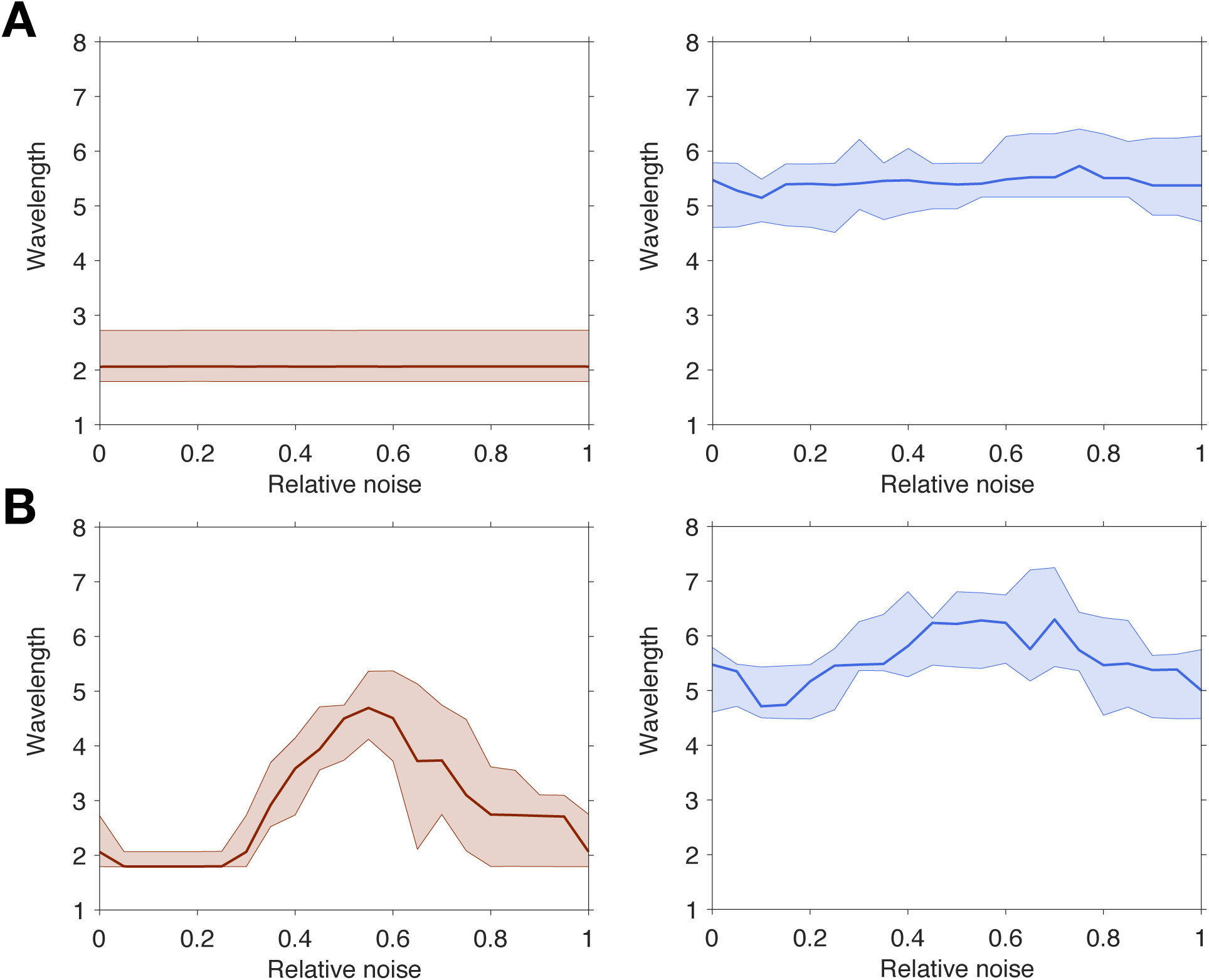
Robustness to noise: The auxin production rate *s*_*a*_ (A) or the PIN1 concentration *P* (B) is spatially and temporally random, with a uniform distribution centered around the value of these parameters without noise. The relative noise amplitude is half the ratio between the width of this interval and the average value of the variable. The wavelength is measured as the average distance between a peak and its nearest neighbor. The prediction of the model with the stress feedback is shown in red (left), while the one with the strain feedback is represented in blue (right). The curves represent the median and the shaded areas the interval between the 15^th^ and 85^th^ percentiles.

### 2.8 Robustness to sharp variations

Even though robustness to noise discriminates between the two models, noise is difficult to manipulate experimentally. Robustness can also be tested by applying a transient shock to the system, i.e. a sharp modification of the external parameters, and by observing whether the tissue returns to its pre-shock state. Experimentally, this is feasible by transient chemical perturbation, such as external auxin application or inhibition of auxin transport, as performed in [18].

We simulated such perturbations as follows. In a tissue at equilibrium, we modified the mean auxin level ⟨*a*⟩ (through the production rate *s*_*a*_, as shown in Fig. 8A) or the concentration of transporters *P*, as shown in Fig. 8B, and let the system reach a new equilibrium. Then we reset the parameter to its initial value and obtain a third equilibrium. We compared the wavelengths of the first and third equilibria. Observing the same wavelength would mean that the system exactly recovers its initial state.

**Fig. 8.**
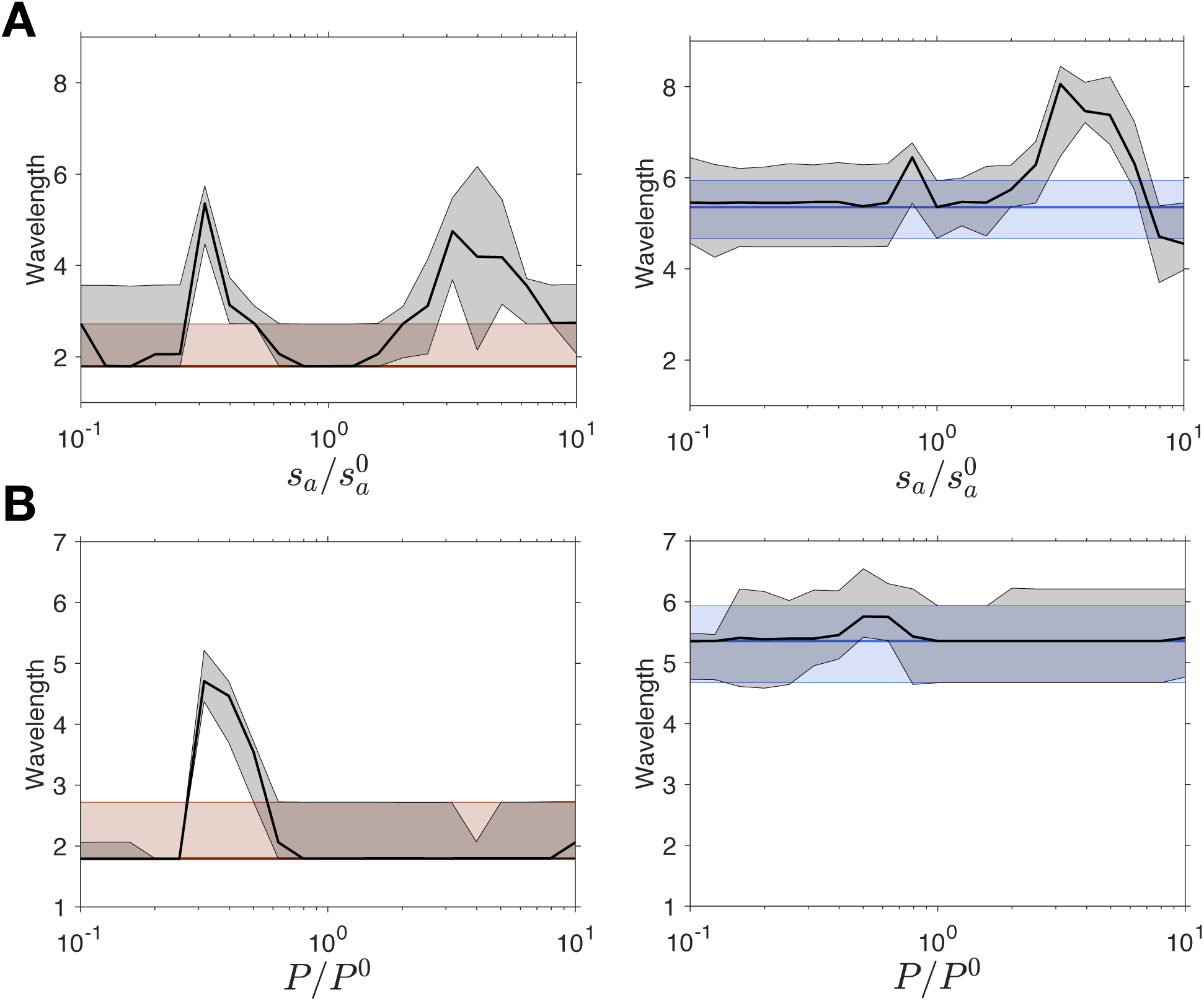
Robustness to sharp variations: The auxin production rate *sa* (A) or the PIN1 concentration *P* (B) is transiently modified over the entire tissue. Once the tissue reaches equilibrium, the original set of parameters is restored. The wavelength is measured at equilibrium before the shock and, and at equilibrium after reseting the original parameters. The wavelength before the shock is plotted in red for the stress feedback (left) and in blue for the strain feedback (right). The wavelength after the recovery is plotted in black. The curves represent the median, the shaded areas the interval between the 15^th^ and 85^th^ percentiles before the shock, and the hatched areas the same percentiles after the resetting.

Fig. 8 shows that large differences in wavelength can be observed between the first and the third equilibria, with both feedbacks. The strain feedback, however, is not as sensitive, compare the left and right columns in Fig. 8. This is qualitatively consistent with its lower sensitivity to noise, documented in the previous subsection.

## 3 Discussion

We developed here a mechanochemical model for auxin patterning in the plant shoot apical meristem. The central question, addressed with our model, is whether the insertion of the efflux facilitators (transporters) in the membrane depends on the strain of the cell walls or by the stress applied to them. To this end, we compared the predictions of the model with available experimental results.

We found that both stress and strain-based feedbacks lead to similar predictions when simulating cell ablation, turgor-induced changes in tissue tension, or a global reduction of the stiffness of the cell walls, generally in agreement with observations. Modeling *PMEI*-overexpressing plants allowed us to discriminate between the two models: Assuming that auxin effect on cell wall stiffness is reduced in such plant lines, we found that only the strain-based feedback directly accounted for observations. Interestingly, organogenesis is abolished in these lines, showing that it is useful to study the system behavior in the absence of patterns. We also compared patterns of auxin and PIN1 concentrations, and show both numerically and analytically that only a strain-sensing mechanism can explain their correlation, as observed in incipient primordia, whereas a stress-sensing mechanism leads to anti-correlated patterns. Note that we assume the amount of transporters per cell to be constant, whereas experiments show that the application of auxin increases PIN1 concentration [18]. Although we cannot rule out that this effect compensates for the observed anti-correlation with a stress-based feedback, a strain-sensing mechanism seems more plausible. Finally, investigating the effect of noise, or of a transient change in chemical parameters, on patterning strengthened our conclusion: Patterns appear more robust with the strain-based feedback.

The model, however, partly failed to reproduce the observations concerning the effect of tissue tension on polarity. Irrespective of the feedback chosen, we found that polarity increased with tension, whereas, experimentally, polarity slightly decreases from isotonic to hypertonic conditions [32]. Again, a possible explanation is that we assumed the amount of transporters per cell to be constant, whereas PIN1 levels decrease in hypertonic conditions; other processes might be triggered when plants react to such osmotic stress. Alternatively, the slight decrease observed could be due to a small experimental bias, and the difference between experiments and our simulations may not be very significant. In addition, the model yielded wavelengths that were smaller for stress- than for strain-sensing. As we have found no explanation for this difference, it is difficult to know whether the model is incomplete or whether this result would also favor strain-sensing because auxin peaks are observed to have a smaller spatial extent than inter-peak distance, which resembles more simulated strain-sensing (Fig. 1C, right). Because of this difference in wavelength, we also explored other values of parameters for the stress-based feedback and found that all conclusions on the comparison with the strain-based feedback hold, except for the conclusions on robustness that are more sensitive to parameter values (supplementary text, Figures S2-S7).

The idealized hexagonal geometry of the cells and the absence of tissue growth are important limitations of the model. The agreement of the analytical linear stability analysis with simulations (supplementary text, Figure S1) makes it likely that cell topology has little effect on our conclusions. Nevetheless, plants respond to mechanical stimuli by altering their growth rates [31]. Increase in auxin levels induce cell growth [7], which induces tissue reorganization, changes in mechanical stress [2], and ultimately feeds back on auxin transport. Although these processes are important in development, mechanical signals are quasi-instantaneous and mostly depend on the current state of the tissue, so that we we do not expect them to significantly our main conclusions.

Altogether, our results favor a feedback based on strain, though many of the experimental configurations are insensitive to whether the mechanical feedback on auxin transport is provided by stress or strain. This raises the question of the underlying molecular mechanisms. Many types of mechanosensors are known [38]. In the case of PIN1, strain could shift the balance between endocytosis and exocytosis, accounting for the strain-based feedback, because osmotic stress affects cell trafficking, in particular through clathrin-mediated endocytosis [53]. The contact between the plasma membrane and the cell wall is needed for PIN1 polarity [5, 10], suggesting that the mechanical state of the cell wall is relevant for polarity. However the role of the cell wall might just be to reduce lateral diffusion of PIN1, helping to maintain the distribution of PIN1 determined by cell trafficking [27].

The question of strain- and stress-sensing was raised in the different, but related, context of plant cortical microtubules orientation by mechanical cues, and the combination of experimental and theoretical approaches suggests that stress-sensing is more likely to be involved [15, 6]. The same conclusion has been reached concerning the actomyosin cortex in the drosophila wing disc [25]. However, the case of isolated animal cells has been debated. Early experiments suggested force-[12] or deformation-sensing [43]. More recent experiments showed that deformation-sensing occurred at low force, while force-sensing occurred at high force [52], though other experiments showed that cells could sense the stiffness of extracellular space [30], consistently with the observation that cells can differentiate according to the stiffness of their environment [3]. Actually, mechanosensing occurs at different scales [38], so that the mechanical variable sensed may depend on the sensing molecule and on the specific function associated with sensing.

In order to disentangle the parameters involved in mechanosensing, it is necessary to combine experimental perturbations of cells or tissues with analytical and computational studies of their behaviors. This is now made possible by the improvement of micromechanical [28] and computational [8] approaches. In this spirit, the present work provides insight on how the interaction between biomechanical and biochemical fields may contribute to the robustness of morphogenesis.

## Model formulation

### Tissue mechanics

We model auxin transport through anticlinal cell walls in the epidermis and thus we neglect the mechanical contribution of other cell walls. We assume that the rest state of the tissue to be a regular hexagonal tiling of the plane and we formulate the problem in terms of a vertex model with periodic boundary conditions. The equilibrium positions of the vertices are obtained by minimizing the mechanical energy of the *N*_*C*_ cells. The contribution to the mechanical energy of the cell wall common to adjacent cells *i* and *j* has a linear density

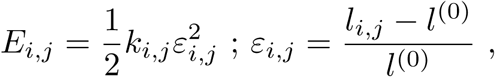

where *k*_*i,j*_ is the stiffness of this wall, *l*_*i,j*_ its length (with an equilibrium value equal to *l*^0^), and *ε* _*i,j*_ its strain. The stress in the anticlinal wall is then given by the derivative of its energy with respect to strain:

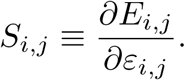

The forces resulting from turgor pressure and tissue curvature are accounted for by external stress with components *σ*_*x*_, *σ*_*y*_. The total energy of the tissue then takes the form

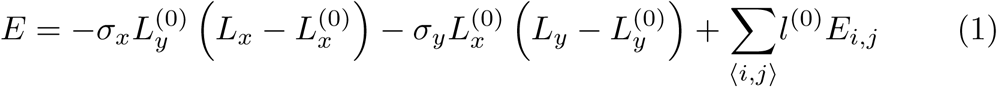

where *L*_*x*_ and *L*_*y*_ are the tissue dimensions along the *x* and *y* directions and 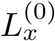 and 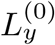 their values without external stress; the sum is over the pairs of neighboring cells ⟨*i, j*⟩. Here we considered only the case of isotropic stress, such that *σ*_*x*_ = *σ*_*y*_ = *σ*.

### Auxin dynamics and coupling with the mechanics

We use the same assumptions as in previous studies [22, 44]. Namely, we only model auxin concentrations in the cytosol. We also assume that PIN1 dynamics occurs with a time scale that is shorter than the time necessary for the transport of auxin through cell walls. The latter is therefore the limiting step. These assumptions allow us to reduce the model to only one chemical equation that describes the auxin concentration, *a*_*i*_, in cell *i*:

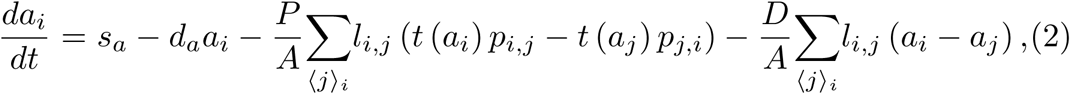

where *s*_*a*_ is auxin synthesis rate, *d*_*a*_ is auxin degradation rate by, and *D* is the diffusion coefficient. ⟨*j*⟩_*i*_ is the set of indices of the 6 cells adjacent to cell *i*. We assume elastic strain to be small, so that the area of each cell is approximated by its rest area *A*. The total amount of PIN1 proteins per cell, 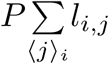, is assumed to be cell-independent; *p*_*i,j*_ is the normalized linear concentration of PIN1 proteins localised at the membrane of cell *i* and facing cell *j* (this PIN1 fraction is responsible for auxin efflux from cell *i* to cell *j*; *P* is used as a unit of linear concentrations). The rate of auxin transport by PIN1 proteins, *t*(*a*), has the following sigmoidal-dependence on auxin concentration:

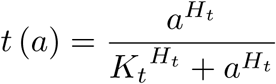

where *K*_*t*_ is a threshold in auxin concentration and *H*_*t*_ the Hill exponent.

Auxin controls tissue mechanics by softening the cell walls (Fig.1). The stiffness of the walls decreases with the amount of auxin in the cells:

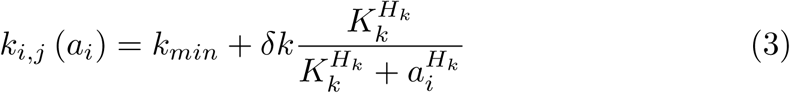

where *k*_*min*_ is the wall stiffness in the absence of auxin, *δk* the variations in stiffness, *K*_*k*_ the auxin threshold and *H*_*k*_ the Hill exponent.

The mechanical effects on the cell also affect auxin dynamics via its transport. The amount of PIN1 transporters in a cell membrane is affected by the strain or the stress, as illustrated in Fig. 1A. The normalized concentration, *p*_*i,j*_, of PIN1 proteins localised at the plasma membrane of cell *i* and facing cell *j* reads

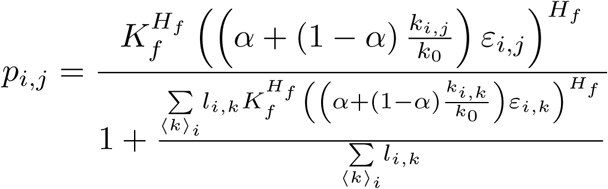

where *α* = 0 for the stress feedback, *α* = 1 for the strain feedback, and *ε*_*i,j*_ and *k*_*i,j*_ stand for wall strain and wall stiffness as described above. The parameter *k*_0_ is set to *k\**, the value of the stiffness in the homogeneous equilibrium state, so that the contributions of stress and strain are of the same order of magnitude in the homogeneous state. *K*_*f*_ sets the amplitude of the insertion function and *H*_*f*_ is the corresponding exponent.

### Observables

Two observables are relevant for the comparison between our simulations and experimental observations, see Fig. 1B [32, 7]. The first is polarity, defined as the ratio of the PIN1 concentrations on the most and least enriched membrane of cell *i*:

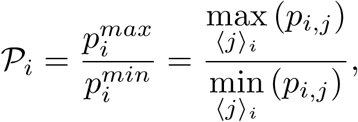

and the second is the ratio of plasma membrane localized PIN1 to the total amount in the cell:

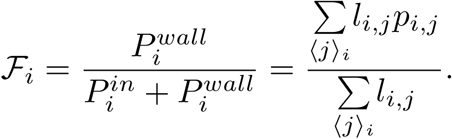

*P* does not appear in this equation because it was used to normalise *p*_*i,j*_. In practice we note 𝒫 and ℱ their distribution of the tissue.

### Implementation

The model was programmed in C. The mechanical energy 1 is minimized thanks to the BFGS algorithm [26], implemented in the NLopt library [21]. The differential equations for the auxin concentration 2 are solved iteratively thanks to the GNU GSL library with a Runge-Kutta (2, 3) method [14]. We assume that the typical time-scale to reach the mechanical equilibrium is much smaller than the time scale for auxin regulation. Consequently, we compute the evolution of the system iteratively, by first computing the evolution of the auxin concentration on a small time-step, and updating the configuration of the tissue by minimizing its mechanical energy. The time-step *dt* = 0.1 for the evolution of auxin dynamics was chosen to limit the number of energy minimization while generating the same results than smaller values. The code is available on Github (github.com/JeanDanielJulien/auxinTransport). Simulations results were analyzed and figures were made with Matlab. Table 1 summarizes the values of the parameters used in the present work.

## Acknowledgements

This work was supported by the DFG SFB 937 “Collective behavior of soft and biological matter”, project A9, to AP, and by the Human Frontier Science Program and the ANR MechInMorph (ANR-12-BSV2-0023) to AB.

## Supplementary material

### Linear stability analysis

The analytical study of the system is performed in the case of isotropic external stress such that σ_*x*_ = σ_*y*_ = σ.

The system is linearised by considering small variations of auxin around the homogeneous stable state *a*\* + *δ*_*i*_ where *a*\* = *s*_*a*_*/d*_*a*_. In the homogeneous stable state we can write the deformation of the walls, their stiffness and the rate of PIN1 insertion as:

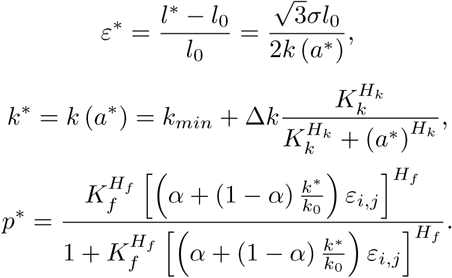

We assume that the local variations in the mechanical tension are small compared with the variations of auxin. Consequently we can write:

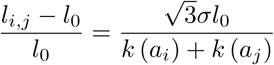

We then linearise the system:

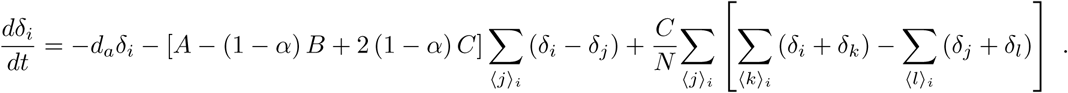

The constants *A, B* and *C* are defined as:

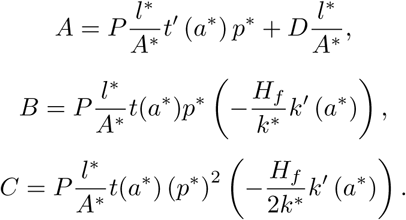

The system can be diagonalised by applying a Fourier transform 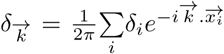, where 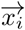 is the position of the cell *i*. Thus we obtain:

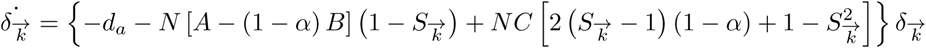

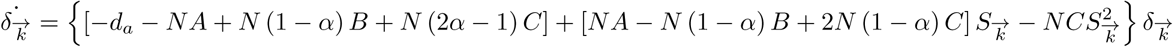

where 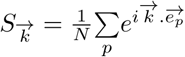 is the form factor of the tissue.

Then we can write the conditions of instability for the two models:

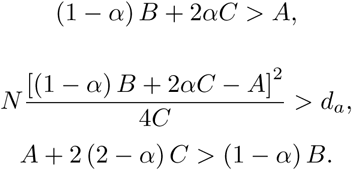

If these conditions are true, we can show that the most unstable wave vectors 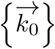 fulfils

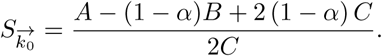

Note that 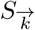 is a decreasing function of 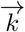, and therefore an increasing function of the wavelength. Since 2*C* = *p*^*^*B< B*, the wavelength for the stress feedback (α = 0) is always smaller than the wavelength for the strain feedback (α = 1), for any set of parameters.

We compared these results with the wavelength measured in simulations for 2-fold changes in the value of each parameter, see Fig. S1. Overall, the linear stability analysis presented here provides a semi-quantitative explanation for the results of our numerical simulations.

### Investigation of different parameters for the stress feedback

The most parsimonious set of parameters, presented in the main manuscript, leads to significanlty different wavelengths for the stress- and strain-sensing mechanisms. In order to evaluate the robustness of the results, we present here the same analysis with 4 different parameter changes of the stress feedback. We first tried modifying *k*_0_, the only parameter that distinguishes the two models. Decreasing *k*_0_ increases the wavelength, but does not allow to reach wavelength of the strain-based feedback. Additionally, it increases the typical fraction of transporters inserted in the membrane *p*^*^ to 1, inconsistent with experimental observations. We present the results for *k*_0_ = *k*_*min*_, close to the wavelength saturation (panels 1 in all figures 2 to 7). We then tried to modify 3 parameters that are not involved in the definition of *p*^*^, and which perturb neither the typical deformation of the cells *ε*\* nor their typical stiffness variations Δ *k/k*_*min*_. The three parameters values (panels 2: *H*_*k*_ = 1.06; panels 3: *D* = 17; panels 4: *P* = 30) were tuned by using the linear stability analysis to match the wavelength of the strain-based feedback. For figures 2 to 7, transporters concentration is regulated by stress. The curves all represent median values and the shaded areas the interval between the 15^th^ and 85^th^ percentiles in a tissue of 3600 cells. Parameters are the same as in Table 1 of the main manuscript, except for one parameter, modified in order to get comparable wavelengths for both feedbacks (1: *k*_0_ = *k*_*min*_; 2: *H*_*k*_ = 1.06; 3: *D* = 17; 4: *P* = 30).

The parameters sets presented here support the conclusion of the main manuscript. In particular, the qualitative differences between the stress-feedback and experiments, namely the absence of internalisation of transporters in simulated PMEI mutants (Fig. 4) and the anti-correlation of auxin and PIN1 patterns (Fig. 5), are valid for all other parameters investigated, consistent with the calculations in the limit of small fluctuations. Results regarding patterns robustness to noise (Fig. 6) and perturbations (Fig. 7) are mitigated, with very different sensitivities of the different parameters sets.

**Figure 1:**
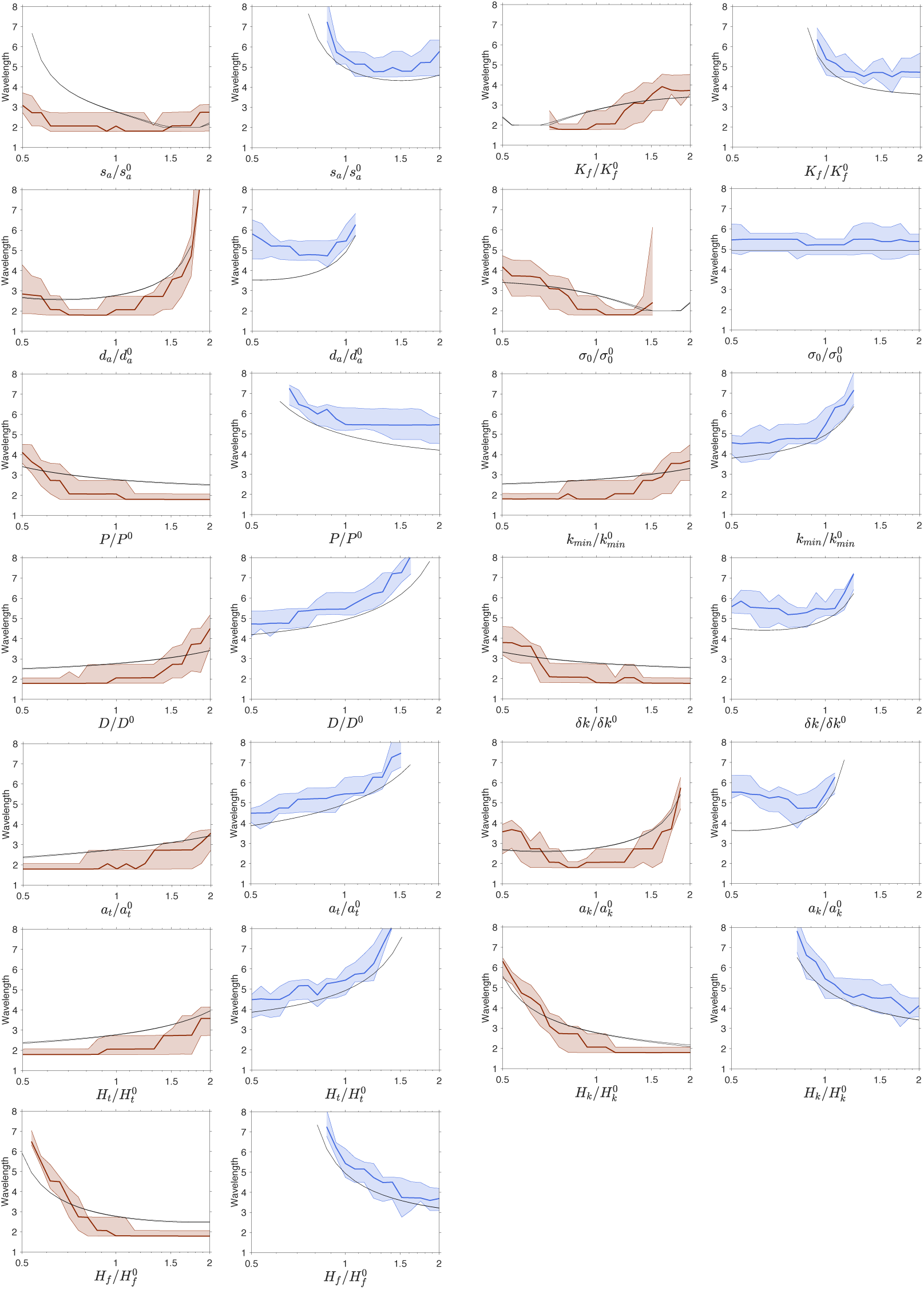
Dependence of the wavelength on each parameter of the model: The wavelength is measured for 2-fold changes in each parameter around the values given in Table 1 of the main text. Results are shown in red for the stress-based feedback, and in blue for the strain-based feedback. The black lines indicate the predictions from the linear stability analysis, in the two directions of the wave vector 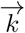 where the prediction is minimal and maximal. Note that these lines almost overlap, except for the smallest wavelengths.

**Figure 2:**
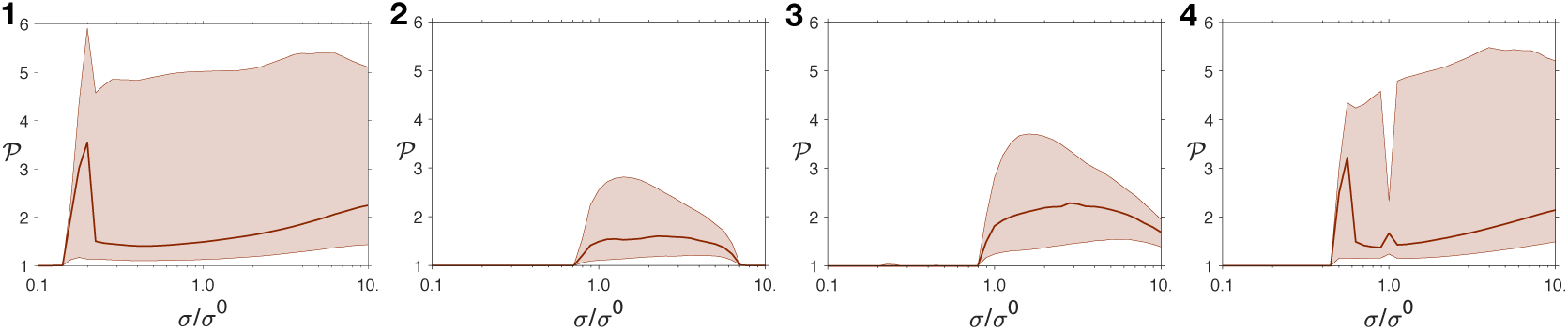
Dependence of polarity on turgor pressure: Starting from σ^(0)^, the pressure is gradually increased or decreased, for pressures ranging from 0.1 × σ^(0)^ and 10 ×σ^(0)^. (1-4) Polarity of transporters for different parameters changes.

**Figure 3:**
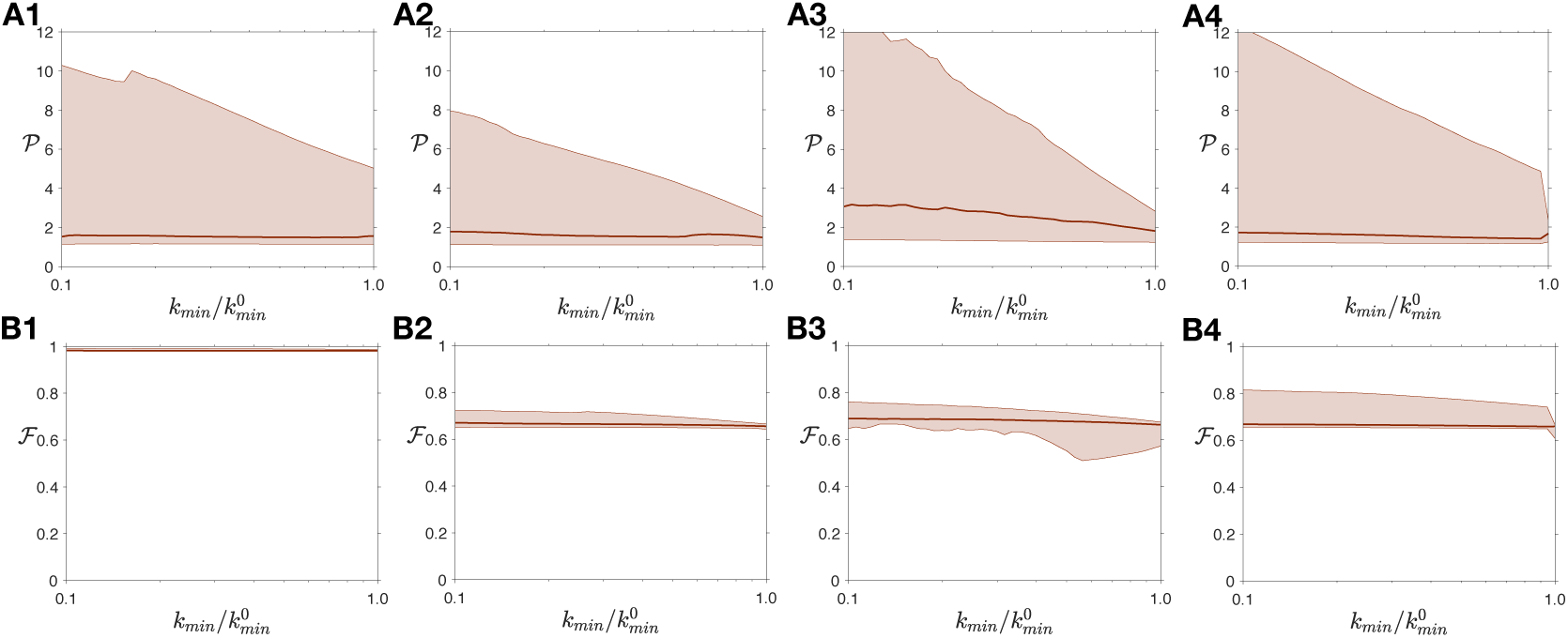
Global softening of the tissue increases polarity: Starting from 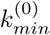, the minimal stiffness of the tissue is gradually decreased, ranging from 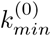 to 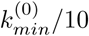. (A1-4) Polarity and (B1-4) fraction of transporters for different parameters changes.

**Figure 4:**
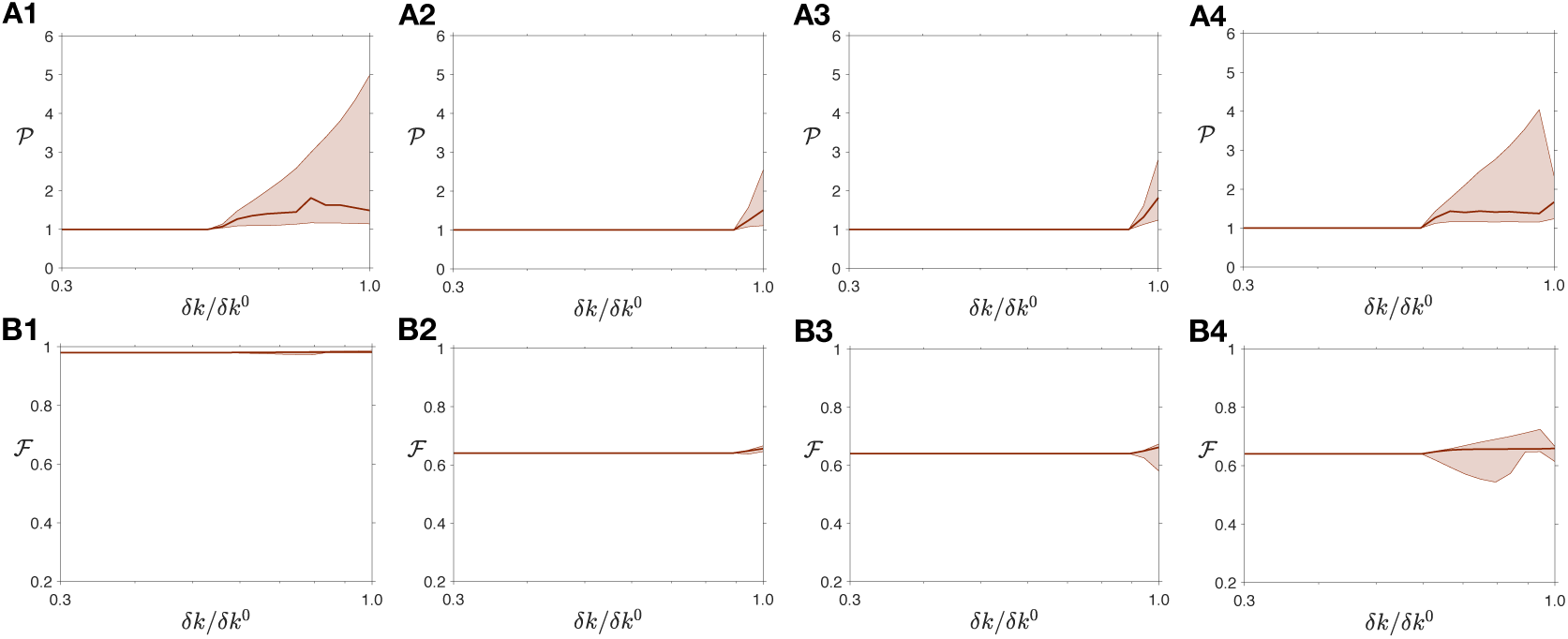
Reducing auxin effects on the cell wall disrupts polarity: Starting from *δk*^(0)^, the amplitude of stiffness variations is gradually decreased, from 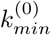 to 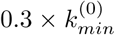. (A1-4) Polarity and (B1-4) fraction of transporters for different parameters changes.

**Figure 5:**
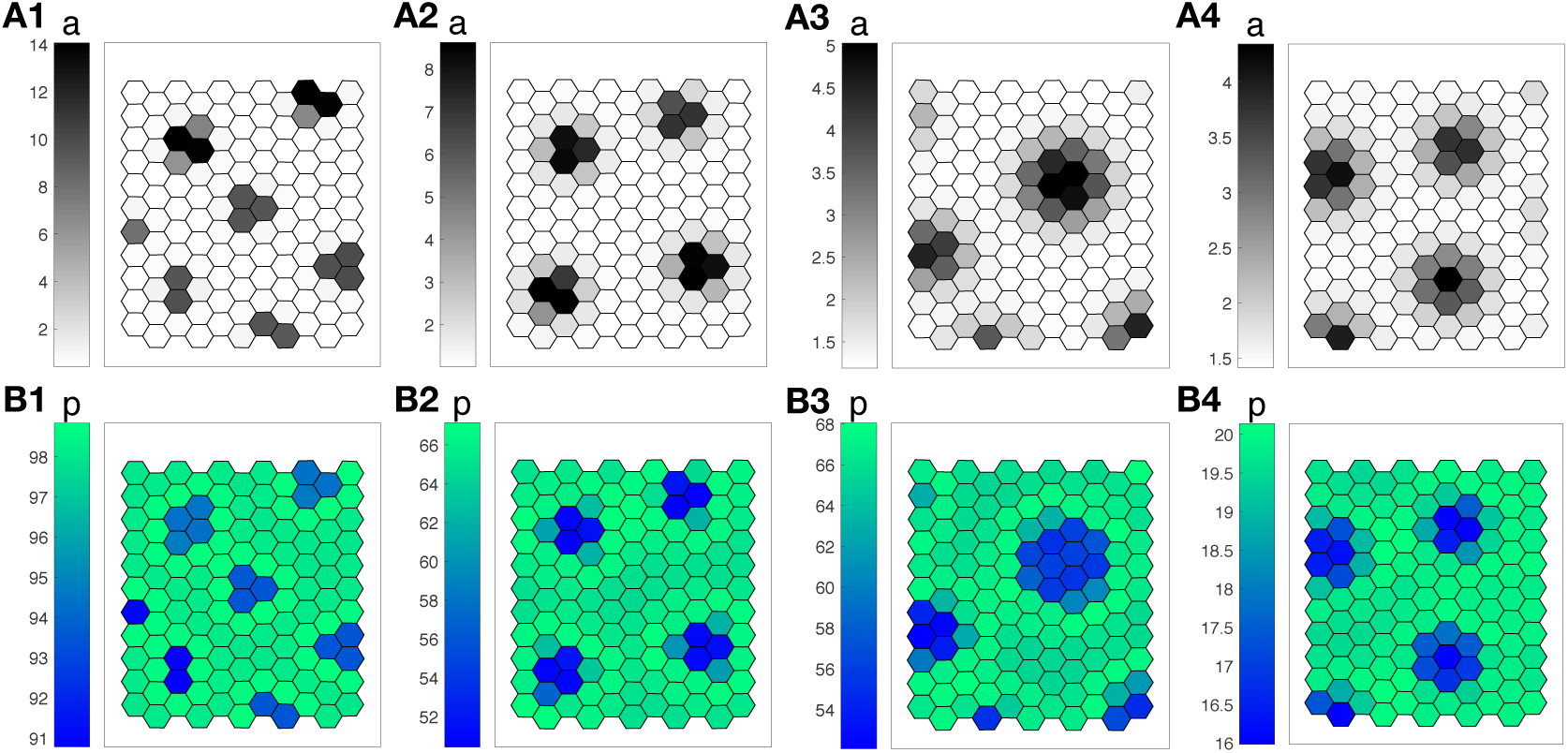
Anticorrelation between auxin and PIN1 concentrations: Examples of pattern predicted by the model, with a stress feedback, for different parameters changes. The cells are colored according to their auxin concentration (gray levels, A1-4), or according to the density of PIN1 transporters, averaged over their walls (blue-green levels, B1-4).

**Figure 6:**
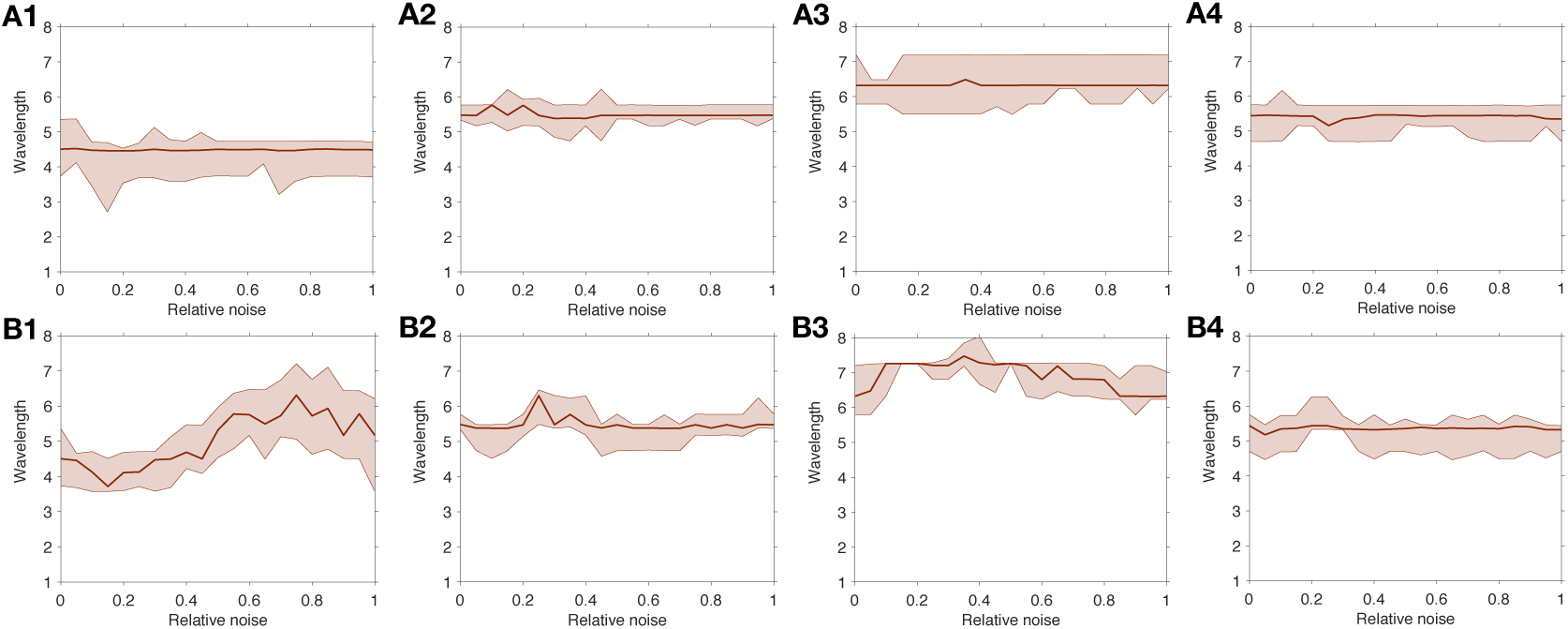
Robustness to noise: The auxin production rate *s*_*a*_ (A) or the PIN1 concentration *P* (B) is spatially and temporally random, with a uniform distribution centered around the value of these parameters without noise. The relative noise amplitude is half the ratio between the width of this interval and the average value of the variable. The wavelength is measured as the average distance between a peak and its nearest neighbor, for different parameters changes (1-4).

**Figure 7:**
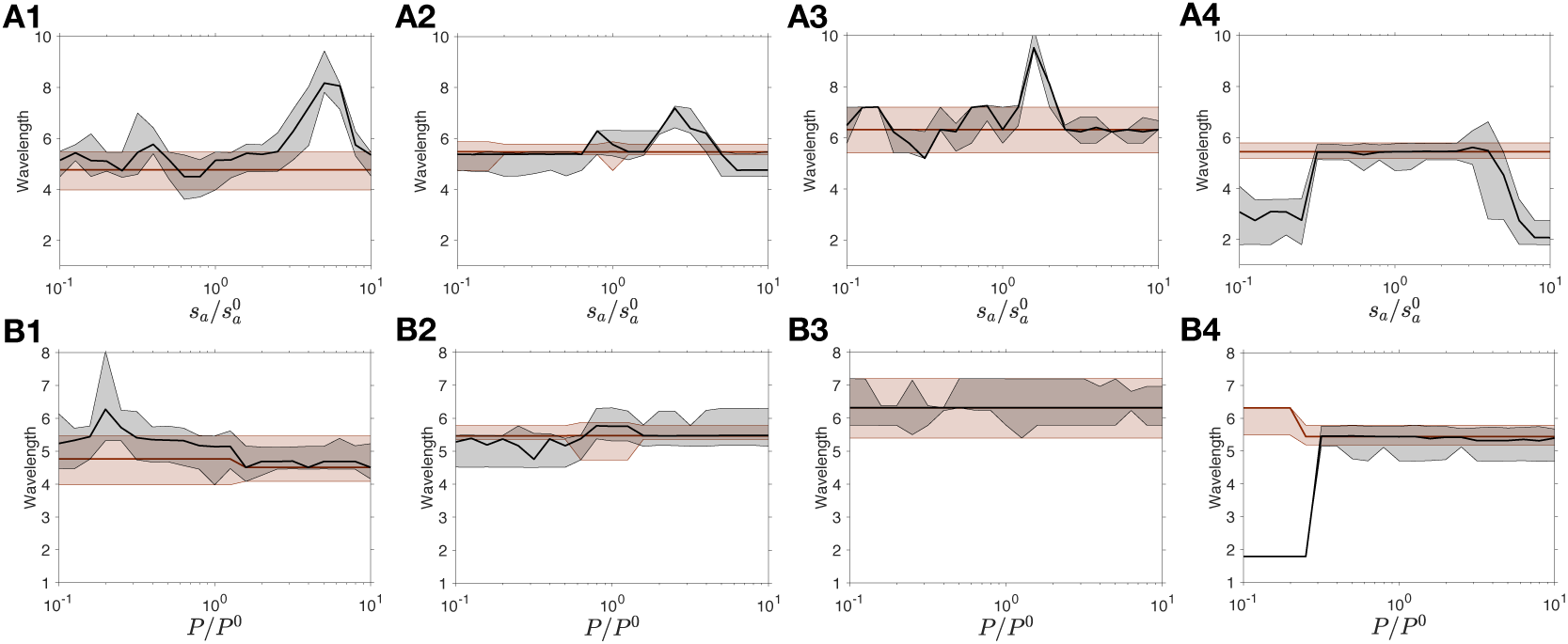
Robustness to sharp variations: The auxin production rate *s*_*a*_ (A) or the PIN1 concentration *P* (B) is transiently modified over the entire tissue. Once the tissue reaches equilibrium, the original set of parameters is restored. The wavelength is measured at equilibrium before the shock and, and at equilibrium after reseting the original parameters. The wavelength before and after the shock is plotted in red and black respectively, for different parameters changes (1-4).

### Change of pattern type with increased pressure Change of pattern type with increased pressure for the stress-feedback

A sharp decrease of the median polarity 𝒫 was predicted with a stress-based feedback when tension is increased above σ*/*σ_0_ = 2.3 (see Fig. 2C). This decrease is due to the change of pattern type. At low tension, the auxin peaks are limited to one or two cells. They become larger as tension increases. This enlargement corresponds to smaller auxin gradients between neighboring cells and accordingly to weaker polarities (see Fig. S8).

### Dependence of the membrane fraction ℱ **on** *δk*

To simulate the effect of pectin methylesterase inhibition, we decrease the parameter *δk*, with the constraint that *k*_*min*_ + *δk* remains constant. By doing so, the dependence of cell wall stiffness with auxin concentration is decreased but the stiffness in the absence of auxin remains the same. The simulations, presented in Fig. 3, show that with the two feedback mechanisms the polarity 𝒫 and the membrane fraction of PIN1 ℱ decrease until the auxin patterns vanish. After the disappearance of the pattern, the simulation results depend on the feedback mechanism. With a stress-based feedback, the fraction ℱ remains constant, whereas with a strain-based feedback it keeps decreasing. These different behaviors can be explained analytically. In the homogeneous state, the membrane fraction ℱ of transporters is:

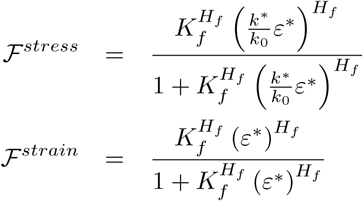

where the star indicates the values in the homogeneous state. *K*^*^ is given by

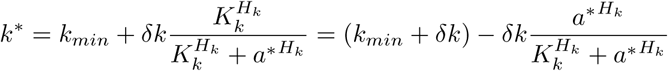

**Figure 8:**
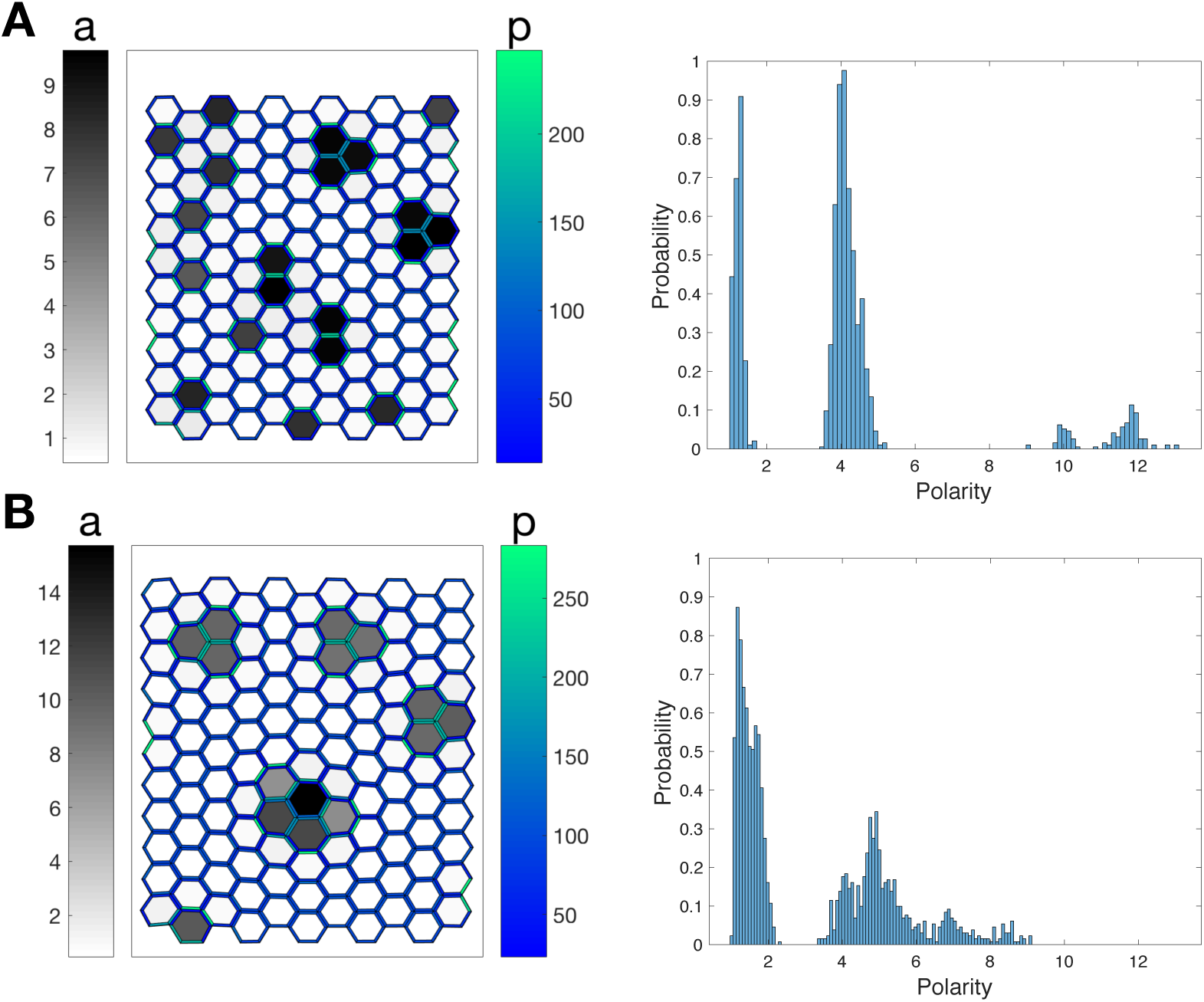
Change in the overall pattern when tension increases: (A) Tissues are represented for σ*/*σ_0_ = 1.41 (left) and σ*/*σ_0_ = 3.55 (right). Each hexagon is a cell that is colored according to its auxin concentration. The two tissues correspond to the parameters marked by black crosses in Fig. 2B. (B) Probability distributions of the polarity 𝒫 computed on the respective tissues.

If we decrease the value of *δk* from an initial value *δk*^0^ with the constraint that *k*_*min*_ + *δk* remains constant, equal to its initial value 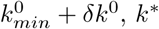 is then:

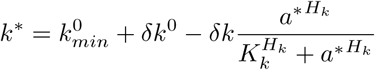

Thus *k*^*^ is a decreasing function of *δk*. The deformation of a regular hexagonal lattice of springs with rest length *l*_0_, stiffness *k*^*^ and under a tension σ is given by 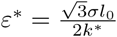.

In conclusion, the fractions are

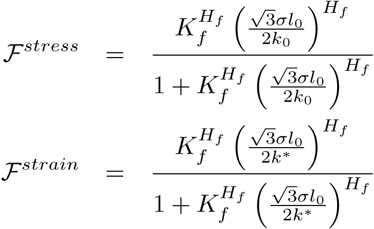

These formulas show that ℱ^*stress*^ does not depend on *δk* whereas ℱ^*strain*^ actually decreases because *k*^*^ increases, The second behavior is consistent with available experimental results [**?**].

### Correlation between auxin and transporters concentrations

We want to study the variation of the mean concentration of transporters in a cell’s membrane,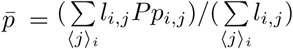, as a function of small fluctuations of auxin *δ*_*i*_ around the homogeneous equilibrium.

Since we assume that the total concentration of transporters is the same in each cell, this is equivalent to the variations of the fraction 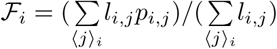.

The change in fraction *δ*ℱ_*i*_ is given by

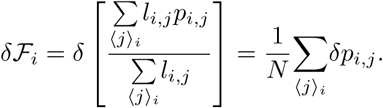

Recall the expression for *p*_*i,j*_:

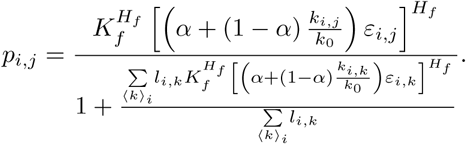

We use the same assumption than for the linear stability analysis to write the fluctuations in strain and wall length as

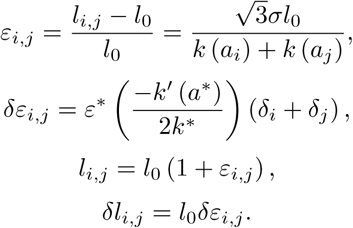

Plugging these equations back in the PIN1 concentration, we obtain:

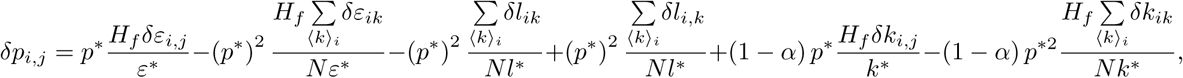

from which we get

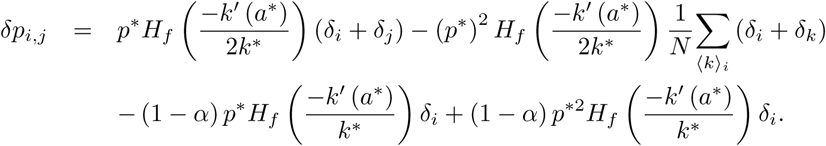

Summing over the neighbours gives the change in fraction *δ*ℱ_*i*_:

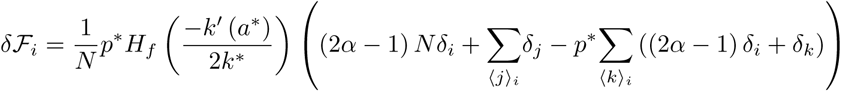

We define the Fourier transform of 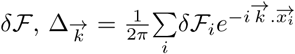, with the Fourier transform of the auxin fluctuations denoted as 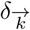, and the form factor of the tissue 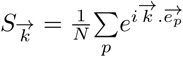.

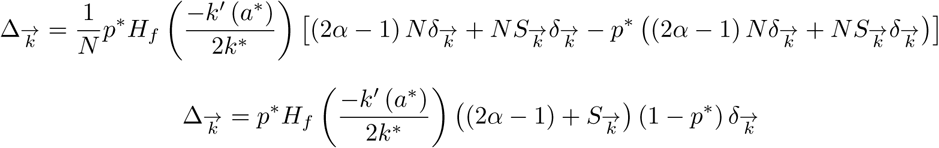

We thus obtain the ratio between transporters and auxin fluctuations, for each feedback mechanism (*α* = 0 corresponds to stress and *α* = 1 to strain):

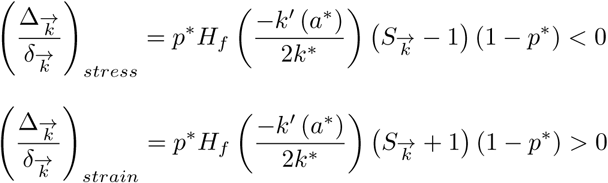

As a conclusion, fluctuations in auxin and PIN1 concentrations have the same sign for the strain feedback, and opposite signs for the stress feedback (we remind that *k*′ *<* 0). This result supports a feedback from mechanical strain, since observations show that both auxin and PIN1 accumulate in incipient primordia. Simulations are in agreement with this result, see Figure 5.

